# Niosome encapsulated doxycycline-hyclate for potentiation of acne therapy: formulation and characterization

**DOI:** 10.1101/2021.09.28.462256

**Authors:** Fatemeh Kashani-Asadi-Jafari, Afra Hadjizadeh

**Affiliations:** Department of Biomedical Engineering, Amirkabir University of Technology, Tehran 159163-4311, Iran

**Keywords:** Niosome, Doxycycline-hyclates, Acne, Antibacterial

## Abstract

Acne is the pilosebaceous units’ disorder. The most important cause of acne is the colonization of bacteria in the follicles. Among antibiotics, doxycycline-hyclate kills a wide range of bacteria. To prevent oral administration’s side effects, overcome the barriers of conventional topical treatment, and improve the therapeutic effectiveness, doxycycline-hyclate was loaded into four niosomal formulations with different percentages of constituents (span 60 and cholesterol) prepared by the thin-film hydration method. Then, one of the four formulations with the most appropriate particle size of 362.88 ± 13.05 nm to target the follicles, percentage of drug entrapment efficiency of 56.3 ± 2.1%, in vitro drug release of 54.93 ± 1.99% after 32 hours, and the lowest permeation of the drug through the Wistar rat skin, was selected. Then, its toxicity on human dermal fibroblasts (HDF) by MTT method after 72 hours, its antibacterial activity against the main acne-causing bacteria via antibiogram test, and its effect on Wistar rat skin drug deposition were measured. Improved cell viability, increased antibacterial activity, and an approximately three-fold increase in drug deposition were the optimal niosomal formulation features relative to the free drug. Overall, this study demonstrated the ability of nano-niosomes containing doxycycline-hyclate to treat skin acne.

## Introduction

Acne vulgaris is an inflammatory disease of the skin pilosebaceous units [1]. The growth and colonization of an anaerobic bacterium called Propionibacterium acne (P. acne) in pilosebaceous units is the leading cause of inflammation in acne disorder [2]. So targeting this microorganism is one of the ways to treat acne disorder. However, systemic antibiotic treatments for acne disorder have limitations such as drug metabolism and low bioavailability. In addition, increasing the dose of the drug in order to overcome these limitations and affect the acne’s pathogenesis causes side effects. Therefore, topical acne treatment is more attractive [3].

The use of conventional topical antibacterial formulations for acne treatment has many side effects for the skin (such as dryness, redness, flaking) because the drug alone cannot target the skin pilosebaceous units (the main center of acne disease). This inability is due to the outermost layer of the skin epidermis, called the stratum corneum, which acts as a powerful barrier to the permeation of external factors into the skin. Therefore, balancing the permeation of a foreign substance through the skin as a therapeutic agent and its deposition in the skin layers at the desired time is essential for treating acne disorder [4].

Vesicular nanocarriers with permeation-enhancing, targeted, and controlled drug delivery have been considered to overcome the side effects mentioned above [1,5]. In addition, nanocarriers increase therapeutic efficacy over conventional formulations due to their targeted delivery and controlled drug delivery characteristics [6,7], so effective results can be achieved by loading drugs with lower concentrations in these nanoparticles. Therefore, bacterial resistance and adverse side effects on the skin are reduced, and patient adaptation is increased [8].

One of the lipid-based nanocarriers is niosomes. They are composed of non-ionic surfactants, usually cholesterol and a hydration medium. Niosomes are made by the self-assembly of non-ionic surfactants [9]. These nanocarriers have a structure similar to liposomes. Inexpensive and affordable raw materials for manufacturing, sterilization, mass production, and higher physical stability are the advantages of niosomes over liposomes [10–12]. Lipid-based nanoparticles form a thin film on the skin, thus preventing trans-epidermal water loss and increasing skin hydration, which improves the therapeutic agent’s permeation through the skin and the deposition of the drug in the local of action relative to the drug without a nanocarrier [13,14].

Doxycycline is a second-generation bacteriostatic tetracycline. Its molecular weight is 512.9 g/mol, and it has a solubility of 50 mg/ml in water. This antibiotic is effective in curing acne by killing and preventing the growth of many gram-positive and negative bacteria, which prevents protein synthesis in them to stop the growth of bacteria [15–17]. However, no optimal nanoparticles have been developed for loading doxycycline-hyclate to treat skin acne, and its oral formulation is usually prescribed to treat acne disorders [18]. Therefore, this drug was loaded into niosomal nanocarriers to reduce side effects, improve therapeutic effects, and overcome barriers to treating acne. In addition, of course, loading this antibiotic into niosomal nanoparticles for ocular delivery and prostate cancer-related infections have been reported [19,20]. However, the synthesis of niosomal vesicles containing doxycycline-hyclate to treat acne with desirable properties and effective against acne-causing bacteria and compatible with skin cells has not yet been reported.

In this study, niosomal nanoparticles with different percentages of cholesterol and span 60 containing doxycycline-hyclate were synthesized. Then the morphological parameters, particle size, zeta potential, the percentage of drug entrapment efficiency, physical stability of samples after 60 days at 4°C, drug release profile in skin simulator environment, effects of synthesized drug-containing niosomal formulations on the in vitro permeation of doxycycline-hyclate through rat skin, the effect of optimal niosomal formulation (N1) on the in vitro drug deposition in rat skin were evaluated. Also, the antibacterial properties of optimal niosomal formulation, the niosomal formulation without the drug, and free drug against two bacteria, P.acne and staphylococcus epidermis (S.epidermis), were performed by the agar well diffusion method. Finally, the cytotoxicity of the optimal niosomal formulation containing the drug and free drug on human dermal fibroblast cells was compared by the MTT method.

## Materials and methods

### Materials

Cholesterol, Chloroform, Span 60, dimethyl sulfoxide (DMSO) were acquired from Merck, Germany. Doxycycline hyclate was purchased from Solarbio, Chine. Tetrazolium salt (MTT) was from Sigma-Aldrich Chemie GmbH, Germany. Dialysis tubing cellulose membrane (12400 molecular weight cut off) was purchased from Sigma-Aldrich, USA. Phosphate buffer saline (PBS) was procured from Zistmavad pharmed, Iran. The human dermis fibroblast cells (HDFCs), Propionibacterium acne (ATCC6919), and Staphylococcus epidermis (ATCC12228) bacteria were obtained from the Iranian Biological Resource Center, Iran.

### Methods

#### Maximum absorption wavelength and standard curve of doxycycline-hyclate

Five samples of doxycycline-hyclate solution in PBS (pH = 7.4) with concentrations of 5, 10, 15, 20, and 40 μg/ml with a volume of 2 ml were prepared to draw the absorption-concentration curve of doxycycline-hyclate. Afterward, the maximum absorption wavelength of the samples was obtained at 276 nm with an ultraviolet-visible spectrometer (Rayleigh, China) (Fig. 1). Finally, the standard absorption-concentration curve was plotted by obtaining the absorption rate of these samples at 276 nm (Fig. 1).

**Fig. 1.**
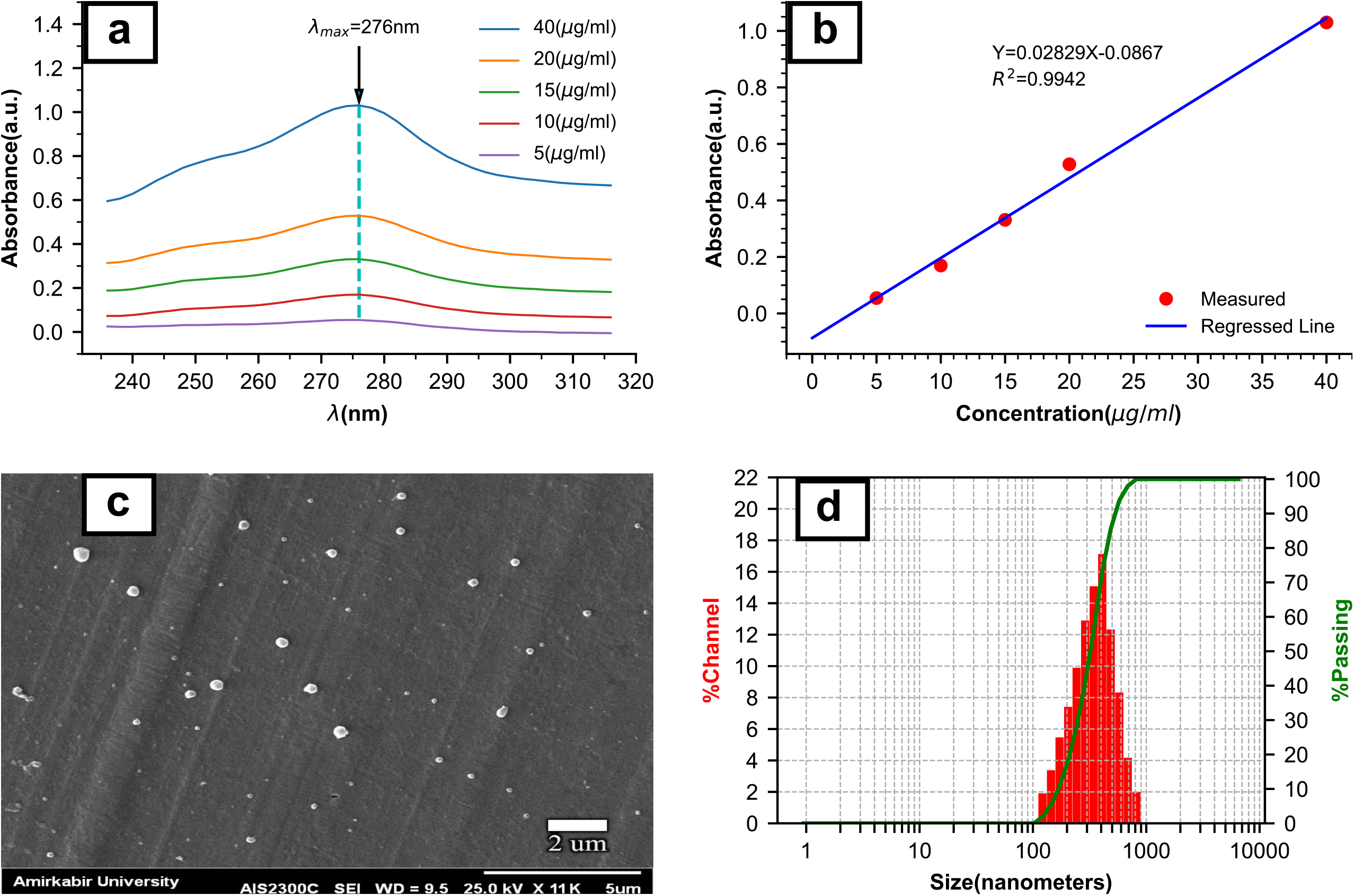
(a) Maximum absorption wavelength of samples at different concentrations of doxycycline-hyclate (5, 10, 15, 20 and 40 μg/ml) that is 276 nm (b) Standard curve of absorption-concentration of doxycycline-hyclate that Y= 0.02829X – 0.0867 (Y is absorption and X is drug concentration) and R^2^ = 0.9942 (R^2^ is the coefficient of determination) (c) Scanning electron microscope image of the N1 formulation (d) The particle size distribution of the N1 formulation.

#### Preparation of vesicles

The thin-film hydration method was used to prepare drug-containing niosomal carriers [21]. First, the appropriate values of span 60 and cholesterol with the total lipid of 700 μmol and molar ratios are shown in Table 1 were dissolved in a round-bottom flask with a capacity of 100 ml, containing 15 ml of chloroform and then transferred to a rotary evaporator (Varghatajhiz, Iran). Next, the chloroform solvent evaporated at 60°C and 120 rpm under reduced pressure. Afterward, 10 μmol of doxycycline were dissolved in 15 ml PBS (pH = 7.4). Next, the thin film inside the balloon was hydrated by the drug solution at 60°C at 80 rpm under normal pressure for 60 minutes, and a milky suspension was obtained at the end. Finally, to reduce the size of the niosomes, the prepared suspensions were sonicated via probe sonicator (Topsonics, Iran) for 10 minutes at 150 watts (8 seconds on, 2 seconds off).

**Table 1.** Molar ratios of span 60 / cholesterol, Diameter size, Zeta potential, PDI, EE%, and LC% for four formulations. Each quantity is displayed as average ± SD for n = 3.

| Formulation | Span60/Chol<br>μM ratio | Diameter size (nm) | Zeta Potential (Mv) | PDI | EE% | LC% |
| --- | --- | --- | --- | --- | --- | --- |
| N1 | 5:5 | 362.88±13.05 | - 24.46±1.39 | 0.17±0.01 | 56.3±2.1 | 19.6±0.7 |
| N2 | 6:4 | 276.25±6.45 | - 29.36±1.22 | 0.23±0.01 | 49.1±1.8 | 16.9 ± 0.6 |
| N3 | 7:3 | 256.17±8.90 | - 38.13±1.60 | 0.26±0.01 | 46.1±1.6 | 15.7±0.5 |
| N4 | 8:2 | 213.76±12.75 | - 44.23±0.85 | 0.29±0.02 | 39.7±1.8 | 13.4±0.6 |

#### Scanning electron microscopy (SEM)

In order to ensure the formation of niosomal vesicles, the morphological characteristics of the prepared samples were investigated with SEM (Seron Technology, South Korea). First, the suspension was diluted with a suitable volume of PBS (pH = 7.4) to reduce the concentration and adhesion of vesicles to each other. Next, the diluted sample was dried at 37°C under vacuum for 24 hours, then was coated with gold via a desktop sputtering system (Emitech, England) and finally subjected to the SEM test.

#### Investigation of particle diameter size, polydispersity index (PDI), and zeta potential

Diameter size, PDI, and zeta potential of niosomes containing doxycycline-hyclate were measured with a dynamic light scattering analyzer (DLS) (Microtrac, Germany). First, each sample was diluted with the appropriate amount of PBS (pH = 7.4), and then the desired properties were measured three times at room temperature, and their average was reported.

#### Entrapment efficiency and drug loading capacity

10 ml of drug-containing niosomal suspension was centrifuged via ultracentrifuge (Phedco, Iran) at 30,000 rpm (100,000 × g) at 25°C for 15 minutes to obtain the percentage of doxycycline entrapment efficiency (EE%) and the loading capacity (LC%). The supernatant containing the free drug was separated from the pellet on the falcon floor in the next step. Then its absorption at 276 nm was measured with an ultraviolet-visible spectrometer. Finally, the concentration of free drug was estimated using the standard absorption-concentration curve of doxycycline-hyclate. With the help of the following equations, EE% and the LC% of doxycycline in niosome vesicles can be obtained [22,23]:

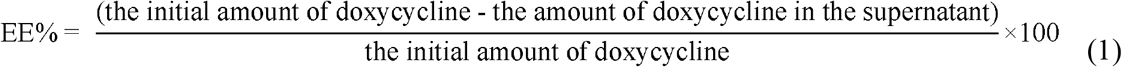

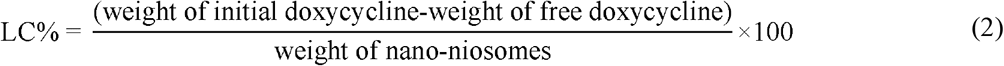

#### Physical stability of niosomes containing doxycycline-hyclate

In order to evaluate the physical stability of niosomal formulations containing doxycycline-hyclate, each sample was placed in a glass vial with a capacity of 8 ml and then sealed with aluminum foil and placed at 4°C for 60 days. After the mentioned period, the values of drug EE%, particle size, and PDI for each sample were calculated and compared with the values of these parameters immediately after synthesis.

#### Evaluation of in vitro drug release

In order to evaluate drug release, each synthesized niosomal formulation containing drug and aqueous doxycycline solution as a control sample was transferred to a dialysis bag (12,400 Mv cut-off) with the equivalent amount of drug. The dialysis bag was then transferred into a beaker with a content of 20 ml PBS (pH = 5.5, Approximate pH of the skin to simulate the skin environment). Then, the beaker was transferred to a heater Stirrer at 37°C and a vortex of 100 rpm to prevent a concentration gradient. Afterward, 1 ml samples were taken from the beaker receptor solution at 0.25, 0.5, 1, 2, 4, 8, 16, 32 hours after the start of the experiment. After each sampling, PBS was added to the beaker at the same temperature and pH to establish the sink conditions. The absorbance of the samples (eight samples) was measured at a wavelength of 276 nm. The amount of drug released at each time was calculated using a standard doxycycline absorption-concentration curve. Thereupon, a graph of drug release percentage versus time was plotted for each sample. Finally, drug release data from each niosomal formulation were analyzed using Zero-order, Higuchi, and Korsmeyer-Peppas kinetic models.

#### In vitro skin permeation

The Franz diffusion cell method was used to investigate the effect of the synthesized niosomal formulations on the doxycycline permeation through the skin [24]. Hairless Wistar rat skin was used to evaluate skin permeation. The vertical Franz used in this study had an effective area of 4.91cm^2^. The skin was hydrated in the receptor solution for 30 minutes, then placed between the donor and receiver chambers so that the dermal area was on the receiver chamber. Each synthesized niosomal formulation and an aqueous drug solution (as a control sample) with the equivalent amount of 750 μg of doxycycline was placed in the donor chamber, and that was sealed. The receiver chamber was filled with 20 ml PBS (pH = 7.4), and the whole device was placed on a heater stirrer at 37°C and under a vortex of 100 rpm. At 0.5, 1, 2, 4, 8, 12, 16, and 24 hours after the start of the experiment, samples of 500 μl were taken from the receiver chamber and quickly replaced with fresh PBS solution at the same temperature in order to establish the sink conditions. The amount of drug permeated through the skin surface unit was estimated by obtaining the absorption rate of the samples at a wavelength of 276 nm with the help of the standard curve of absorption-concentration of doxycycline. In addition, the flux values at 24 hours and the enhancement ratio (ER) (the flux ratio of each formulation to the control sample) were calculated.

#### Skin deposition assessment

This study was performed to evaluate the effect of niosomal formulation on the amount of doxycycline-hyclate deposited in the viable epidermal and dermal layers. For this purpose, after in vitro skin permeation evaluation, the skin (in contact with N1 formulation and the control sample) was removed from the Franz. First, its surface was cleaned with cotton soaked in PBS. Next, to remove the stratum corneum, the skin was placed on a flat surface, and nine tapes with scotch tape (scotch magic 3m 810) were applied [25]. When the stratum corneum was utterly detached, and the skin surface was shiny, the skin was cut into small pieces, then placed in methanol, afterward ultrasonicated for 30 minutes, and then centrifuged. Finally, after passing the 0.22 μm syringe filter, the supernatant was subjected to spectrophotometric analysis at 276 nm to determine the amount of drug.

#### Cytotoxicity evaluation

Human dermal fibroblasts (HDF) and dimethyl-thiazole diphenyltetrazolium bromide (MTT) assay were used to evaluate and compare the degree of cytotoxicity induced by niosomal formulation (N1) and an aqueous drug solution as a control sample [26]. In summary, HDF cells were placed in a 96-cell plate with a density of 1×10^4^ cells with 100 μl of Dulbecco’s Modified Eagle Medium (DMEM) containing 10% fetal bovine serum (FBS) in each well. They were then incubated at 37°C for 24 hours to ensure that the cells were attached to the floor. After this time, 20 μl of N1 formulation and the aqueous doxycycline solution (as a control sample) at drug concentrations ranging from 0.4 to 0.225 μg/ml were placed in certain wells in direct contact with HDF cells for 72 hours. The culture medium in each well was then replaced with 150 μl of diluted MTT with culture medium (0.5 mg/ml) and incubated for 3 hours at 37 ° C. The MTT solution was then removed from the wells, and 100 μl of dimethyl sulfoxide was added to each well to dissolve the formed formazan crystals, resulting from viable cells’ metabolic activity, and shaken for 10 minutes. The optical density of each sample was measured at 570 nm with an ELISA reader (Awareness Technology, USA). Due to the light sensitivity of the MTT assay reagents, this process was performed in a dark environment. The concentration at which cell survival was 50% (IC50) was obtained by linear regression with Microsoft Excel software. The cell viability percentage of each well was calculated using the following equation [27]:

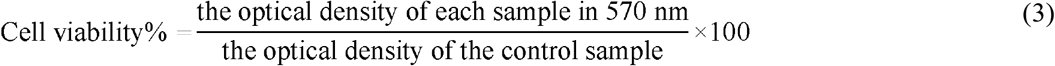

#### Antibacterial activity

To compare the antibacterial activity of the niosomal formulation (N1), an aqueous doxycycline solution, and a drug-free niosome sample against the major acne-causing bacteria, namely P.acne and S.epidermis (an anaerobic bacterium present in acne-causing lesions) [28], antibiogram test and Mueller-Hinton agar medium were used. First, the two bacteria suspensions were cultured using a cotton swab on Muller-Hinton agar, then 10 μl of niosomal formulation (N1), aqueous doxycycline solution, and drug-free niosomal sample were placed on the medium surface and was incubated for 24 hours at 37°C. Finally, the diameter of the growth inhibition zone was measured using a caliper.

#### Statistical analysis

In our study, the data were reported as mean ± standard deviation for n = 3 (n: the number of experiments). Data analysis was performed using SPSS software version 26. Student t-test was used to show statistically significant differences, and the P-value of ≤ 0.05 was evaluated as the lowest significance level.

## Results and discussion

### Preparation of niosomes containing doxycycline-hyclate by thin-film hydration method

According to recent studies, the thin film hydration synthesis method is a suitable and straightforward method for synthesizing niosomes [29]. Therefore, in this study, the same method was used to prepare niosomes containing doxycycline. High EE% and high LC% of the drug in niosomal carriers lead to more favorable therapeutic effects [30]. Therefore, Span 60 was selected due to having the highest phase transition temperature among the non-ionic surfactants used to synthesize niosomes (Tc = 53°C), suitable CPP, and HLB values [31]. By decreasing the chain length of the alkyl in surfactant and increasing the number of unsaturated bonds in it, the Tc decreases, and the permeability of the vesicular bilayer increases. As a result, the probability of leakage of the hydrophilic drugs into the suspension medium increase, and the EE% decreases [11]. In addition, as the HLB increases, the size of the niosomes decreases. This phenomenon could occur due to increased hydrophobicity and consequently decreased surface free energy [32].

### SEM

The morphology of the N1 sample was examined via SEM. As is shown in Fig. 1, the niosomes had an almost spherical structure with a smooth and no adhesion surface.

### Investigation of particle diameter size, polydispersity index (PDI), and zeta potential

As is shown in Table 1, the mean PDI of the samples varied from 0.17 ± 0.01 to 0.29 ± 0.02. A PDI less than 0.1 indicates the desired homogeneity and a narrow peak of the particle size distribution, and a PDI greater than 0.3, indicating a higher heterogeneity [33]. Thus, the obtained PDIs for the prepared niosomal formulations were in the desired range in terms of homogeneity. The particle size distribution of the N1 formulation is shown in Fig. 1, and the average particle diameter size for all formulations is shown in Table 1. It can be seen that as the cholesterol concentration increased, the size of the niosomes increased.

The size of the niosomal vesicles is essential for targeting the follicles in the treatment of acne, which should reduce the transdermal absorption of the drug and increase the deposition of the drug in the skin layers. The best size for nanoparticles to target follicles is in the range of 400 to 700 nm [34]. Therefore, the N1 sample with a particle size of 362.88 ± 13.05 nm was more suitable than other formulations for treating acne due to its proximity to this particle size range. In the structure of niosomes, the hydrophilic head group of cholesterol forms a hydrogen interaction with the oxygen ester group of the span 60. Moreover, its steroid portion is positioned in the direction of the alkyl chains. In this way, the structure of the bilayer membrane is strengthened [35]. On the other hand, in this study, the probe sonication technique was used to reduce the size of drug-containing niosomes, so formulations with higher cholesterol levels were less affected by this size reduction process due to the increased stiffness membranes [36]. Some studies have confirmed the effect of increasing cholesterol concentration on increasing particle size [37,38].

### Characterization of entrapment efficiency and loading capacity

The EE% and LC% of the drug in the prepared niosomal formulations are shown in Table 1. The N1 sample had the highest EE% and LC% of the drug, and this sample was more desirable from this point of view than other prepared niosomal formulations.

The drug EE% increased with increasing cholesterol concentration. It can be stated that the presence of cholesterol in a niosomal formulation abolished the gel-liquid phase transition temperature [39]. In addition, it tightened the structure of the niosomal bilayer membrane. As a result, the leakage of hydrophilic compounds was reduced to the environment outside the niosome [36]. Probably, for this reason, by increasing the cholesterol concentration in the prepared niosomal formulations, the drug’s EE% and LC% increased. The increased hydrophilic drugs’ EE% in niosomal formulations with increasing cholesterol percentage has also been reported in recent studies [40,41].

### Zeta potential analysis

As is shown in Table 1, the zeta potential of the prepared formulations was in the range of −24.46 ± 1.39 to −44.23 ± 0.85 mV.

The zeta potential of drug-containing niosomal formulations is essential to target and improve follicular permeation and deposition [42]. Pilosebaceous units are the site of acne pathogenesis. These units are located in the skin with positive zeta potential [43]. Therefore, it seems that the negative values of the zeta potential of the formulations prepared in this study facilitated their interaction with pilosebaceous units. Furthermore, the zeta potential values of the prepared formulations decreased with increasing cholesterol concentration, which agrees with some of the recent studies [20,44].

### Stability analysis

As is shown in Fig. 2, formulations with lower cholesterol concentrations were less stable than formulations with higher cholesterol concentrations. The color of the samples was also examined, none of which changed color or blurred. The EE% also changed in the range of 8.9% to 13.9% of the amount of drug encapsulated for the samples. However, changes in PDI and size after the mentioned period were not noticeable for any samples, and all samples can be considered stable.

**Fig. 2.**
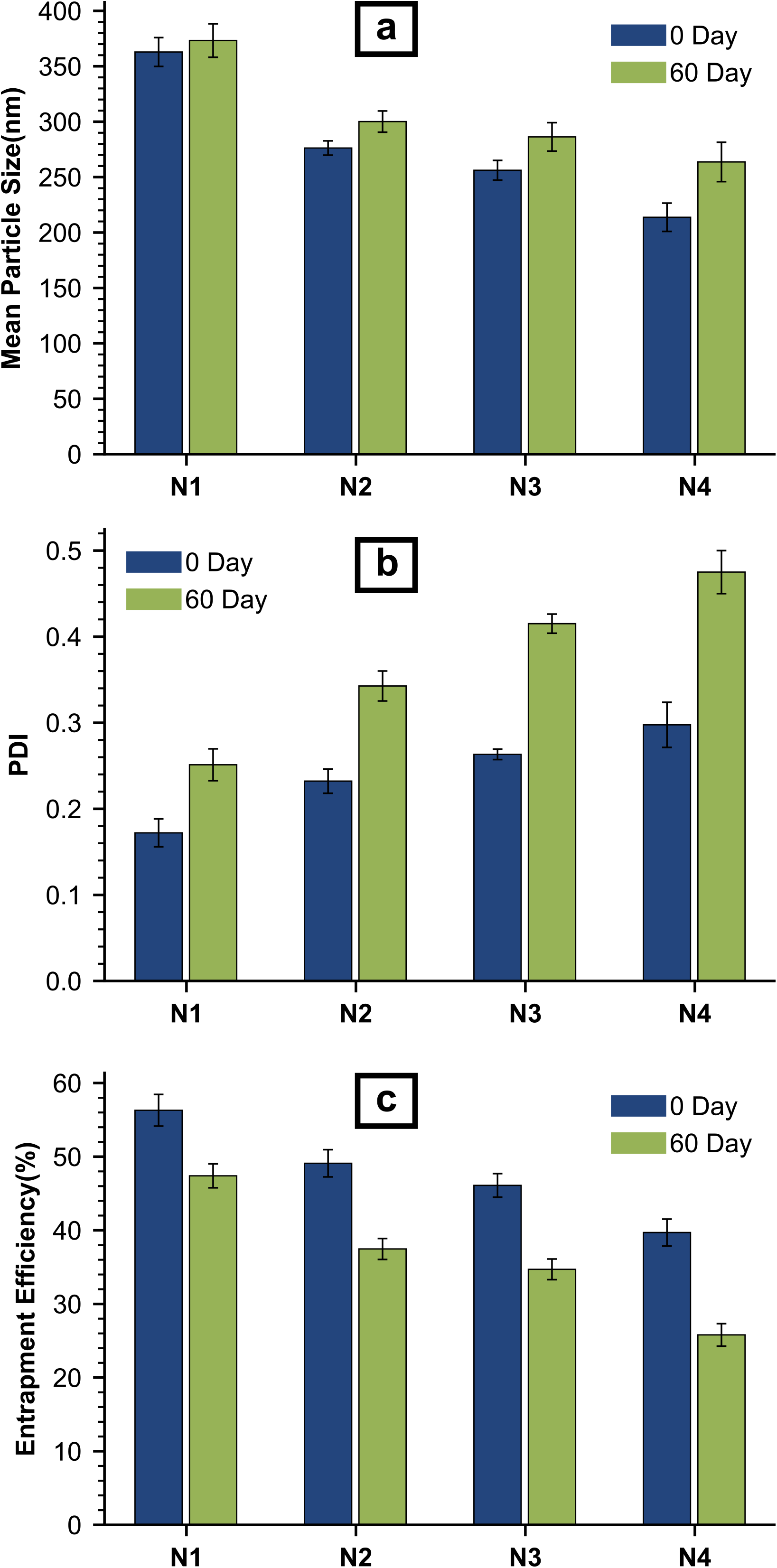
Particle size changes of niosomal formulations after 60 days (b) PDI changes of niosomal formulations after 60 days (c) EE% changes of niosomal formulations after 60 days.

Regarding stability, it can be said that Increasing the size of nanoparticles over time due to their fusion and aggregation can threaten the stability of the formulations [45]. It has also been reported that the stability of formulations is related to the amount of zeta potential. As the zeta potential increases, the repulsive force between the nanoparticles and the stability of the formulations increases [46]. Although in the prepared samples, the zeta potential increased with decreasing cholesterol concentration. However, samples with higher cholesterol concentrations showed more stability. Thus, in addition to repulsive forces between particles, the rate of water absorption and subsequent rupture of niosomes over time also affected the stability of niosomal formulations [47].

### In vitro drug release

As is shown in Fig. 3, the drug release from the aqueous doxycycline solution to the environment after 4 hours was approximately 100%. The percentage of drug release from niosomal formulations after 4 hours was in the range of 29.11 ± 3.74% to 49.74 ± 1.461%, and the lowest drug release occurred in the N1 sample. After 32 hours, the amount of doxycycline released from the niosomal formulations ranged from 54.93 ± 1.99% to 79.87 ± 2.31%. Also, the mechanism of drug release from the formulations, as is shown in Table 2, was most consistent with the Korsmeyer-Peppas model (R^2^ ≥ 97%). At N1formulation, Ficken diffusion (n ≤ 0.43), and at N2, N3, and N4 formulations, a non-Ficken diffusion mechanism occurred (0.43 ≤ n ≤ 1) [48–50].

**Fig. 3.**
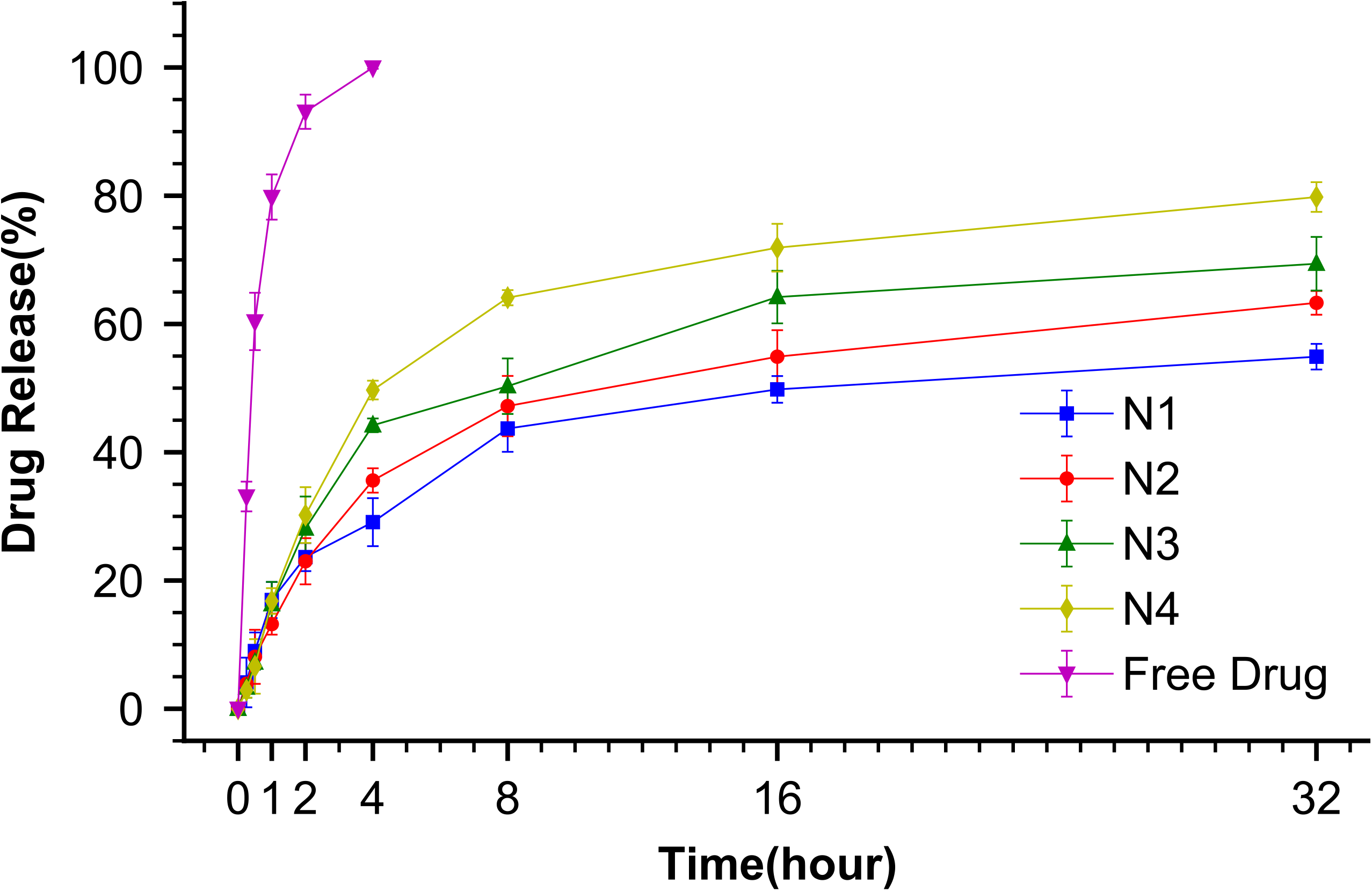
Diagram of percentage doxycycline-hyclate release from four niosomal formulations and aqueous drug solution as a control sample into PBS medium (pH = 5.5).

**Table 2.** Kinetics models for drug release from four prepared niosomal formulations.

| Formulations | Doxycycline release kinetics |  |  |  |  |  |  |
| --- | --- | --- | --- | --- | --- | --- | --- |
| | $M_t/M_{\infty} = k_d t$ | | $M_t/M_{\infty} = k_d t^{0.5}$ | | $M_t/M_{\infty} = k_d t^n$ | | |
|  | k | R<sup>2</sup> | k | R<sup>2</sup> | k | n | R<sup>2</sup> |
| N1 | 2.262 | 0.841 | 11.770 | 0.956 | 17.252 | 0.360 | 0.977 |
| N2 | 2.554 | 0.859 | 13.110 | 0.964 | 15.839 | 0.475 | 0.983 |
| N3 | 2.859 | 0.836 | 14.789 | 0.951 | 17.024 | 0.561 | 0.976 |
| N4 | 3.293 | 0.827 | 17.025 | 0.945 | 15.664 | 0.846 | 0.996 |

Considering the result of drug release, it is evident that niosomal formulations provided sustained drug release. It is also observed that the release of doxycycline from the niosome was biphasic. In the first phase, there was a rapid drug release. This phase was probably related to drugs located on the outer surface of the niosomes, and their release did not require much diffusion. In the next phase, a gradual release of the drug was observed. This phase was probably related to drugs trapped in the center of the niosomal vesicles, the release of which required time to diffuse through the bilayer structure of the niosome [51]. This biphasic release has also been reported in some studies in the past [52,53]. As the concentration of cholesterol in the prepared niosomal formulations increased, the drug release rate decreased. The cause of this phenomenon may have been the removal of the peak phase transition temperature of the niosomal bilayer structure by cholesterol [36].

### In vitro skin permeation

The amount of drug permeation from the dialysis bag could not indicate the amount of drug permeation through the skin when applying the formulations on the skin. That could be due to the skin thickness, the skin appendages, the skin composition, and the interaction between the skin and the niosomal vesicles [54]. Therefore, to investigate the effect of the prepared niosomal formulations on the drug permeation rate more precisely, a vertical Franz and rat skin were used. Wistar rat skin is about eleven times more permeable than human skin. Therefore, it is expected that the transdermal drug absorption in human skin is lower than in rat skin when topical drug formulations are applied [55].

The prepared niosomal formulations enhanced the cumulative drug amount permeated per unit area through skin up to 24 hours after the start of the experiment compared to the control sample. The graph of the cumulative drug amount permeated per unit area through rat skin versus time for the prepared formulations and the control sample is shown in Fig. 4. In addition, flux values at 24 hours and ER for all formulations are also shown in Table 3. ER was in the range of 2.40 to 3.91, and the lowest ER value was related to the N1 sample. On the other hand, flux values for all formulations were significantly higher than the control sample, which showed improved drug permeation through the skin due to applying niosomal formulations.

**Fig. 4.**
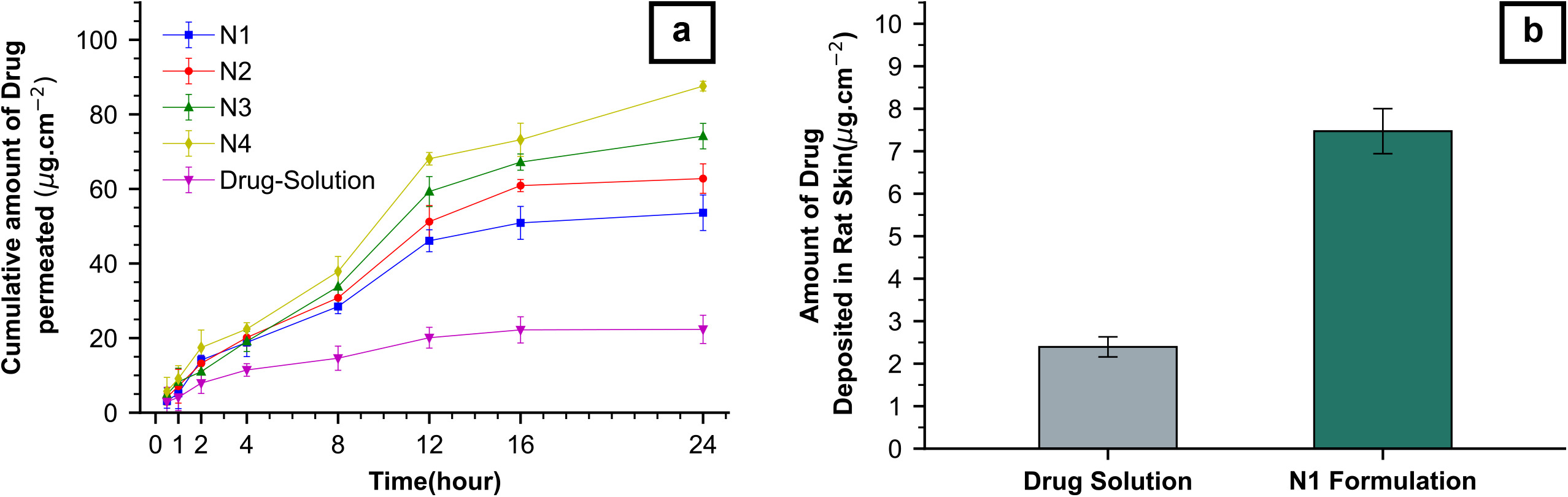
(a) Permeability profile of four niosomal formulations and aqueous doxycycline-hyclate solution as a control sample (b) Comparison of the amount of drug deposition between N1 formulation and aqueous drug solution as a control sample in the rat skin (dermis and viable epidermis).

**Table 3.**
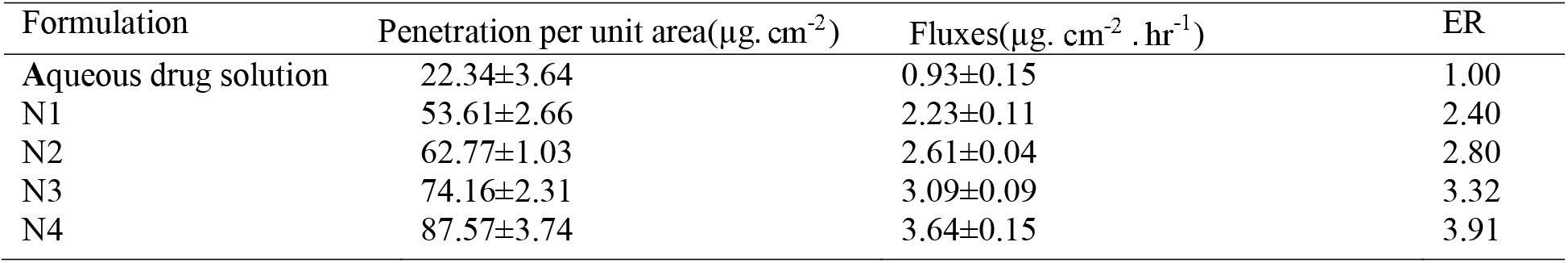
The fluxes (μg. cm^-2^ hr^-1^), drug permeation per unit area (μg cm^-2^) and the enhancement ratio (ER) for four niosomal formulations. Each quantity is displayed as average ± SD for n = 3.

Niosomal vesicles modify the stratum corneum structure by reducing the amount of transepidermal water loss. In addition, they increase the hydration of this layer by weakening the intercellular lipid structure in it. Surfactants in niosomal formulations may also interact with lipids in the stratum corneum to form a drug source on the skin surface with a high concentration gradient. This source of the drug is considered to be the driving force for drug permeation [51]. As the cholesterol concentration in the niosomal formulations increased, the cumulative drug amount permeated per unit area of skin, flux, and ER decreased. The cause of this phenomenon could be increasing the particle size of niosomes and decreasing the concentration of non-ionic surfactants as permeation enhancers. In some recent studies, non-ionic surfactants have been described as permeation enhancers [56,57].

Therefore, this section can conclude that the N1 sample showed lower transdermal delivery than the other formulations. So, due to the appropriate particle size, highest EE% and LC% of doxycycline, optimal physical stability, and the slowest drug release, the N1 formulation was selected for further studies.

### Skin deposition

Fig. 4 shows the amount of drug deposition in rat skin when the niosomal formulation (N1) and the control sample were applied in the equivalent amount of the drug at 24 hours. The amount of drug deposition in the skin (viable epidermis and dermis) when the N1 formulation was applied was 7.47 ± 2.24 μg/cm^2^ that is approximately three times higher than the control sample (Table 4).

**Table 4.** Comparison of the amount of drug deposition between the aqueous drug solution and N1 formulation in rat skin.

| Formulation | Amount of drug deposition in rat skin ($\mu\text{g.cm}^{-2}$) |
| --- | --- |
| Aqueous drug solution | 2.39±1.18 |
| N1 formulation | 7.47±2.24 |

According to the results, increased deposition was probably due to the fusion and absorption of the niosomal vesicles to the skin surface [56]. Another cause of increased deposition could be attributed to the non-ionic surfactants present in the bilayer structure of the niosomes, which altered the arrangement of intracellular lipids in the skin and reduced their crystallinity, thereby increasing skin deposition [58].

### Evaluation of cytotoxicity

As is shown in Fig. 5, the optimal niosomal formulation (N1) at the highest drug concentration (0.225 μg/ml) showed approximately 80.1 ± 6 % cell viability after 72 hours of incubation. On the other hand, aqueous doxycycline solution with similar drug concentration and similar incubation time showed 66.2 ± 4.4 % cell viability. Also, the IC50 values were calculated to be 629.15 μg/ml for the aqueous drug solution and 364.59 μg/ml for the N1 formulation.

**Fig. 5.**
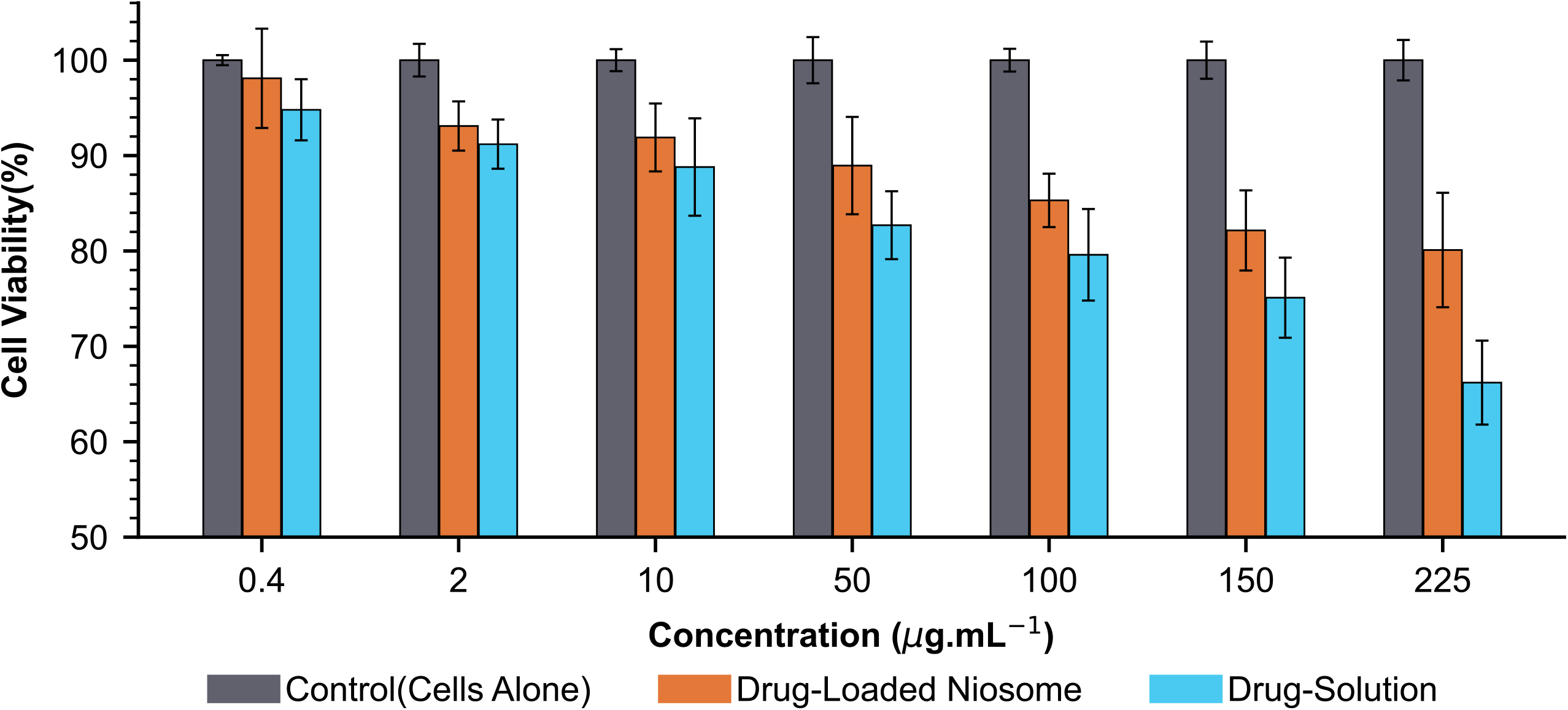
Evaluation of HDF cell viability by applying N1 formulation and aqueous drug solution.

In evaluating cytotoxicity, direct contact of doxycycline with the cells was prevented by loading doxycycline into niosomal vesicles. In addition, the niosomal structure reduced the drug toxicity to HDF cells, probably because the biocompatible and non-toxic components of the niosomal vesicles came in direct contact with the cells, and the drug was gradually released from these vesicles [59,60]. Thus, niosomal nanocarriers could target the drug with similar concentrations with fewer side effects than the free drug. In addition, cell viability decreased with increasing drug concentration if both formulations (N1 and control sample) were applied. This increase in drug-dependent cytotoxicity has also been reported in some recent studies [19,27].

### Antibacterial activity

The results of the antibacterial activity test are shown in Fig. 6. The mean diameter of the growth inhibition zone for the optimal niosomal formulation (N1), aqueous doxycycline solution, and niosome without the drug are given in Table 5. The drug-free niosomal carrier showed little antibacterial properties compared to the control sample against both bacteria. However, the antibacterial activity of the N1 formulation, especially against S.epidermis, was significantly higher than that of aqueous doxycycline. These observations could indicate the potential of niosomal nanocarriers to increase the antibacterial activity of free doxycycline.

**Fig. 6.**
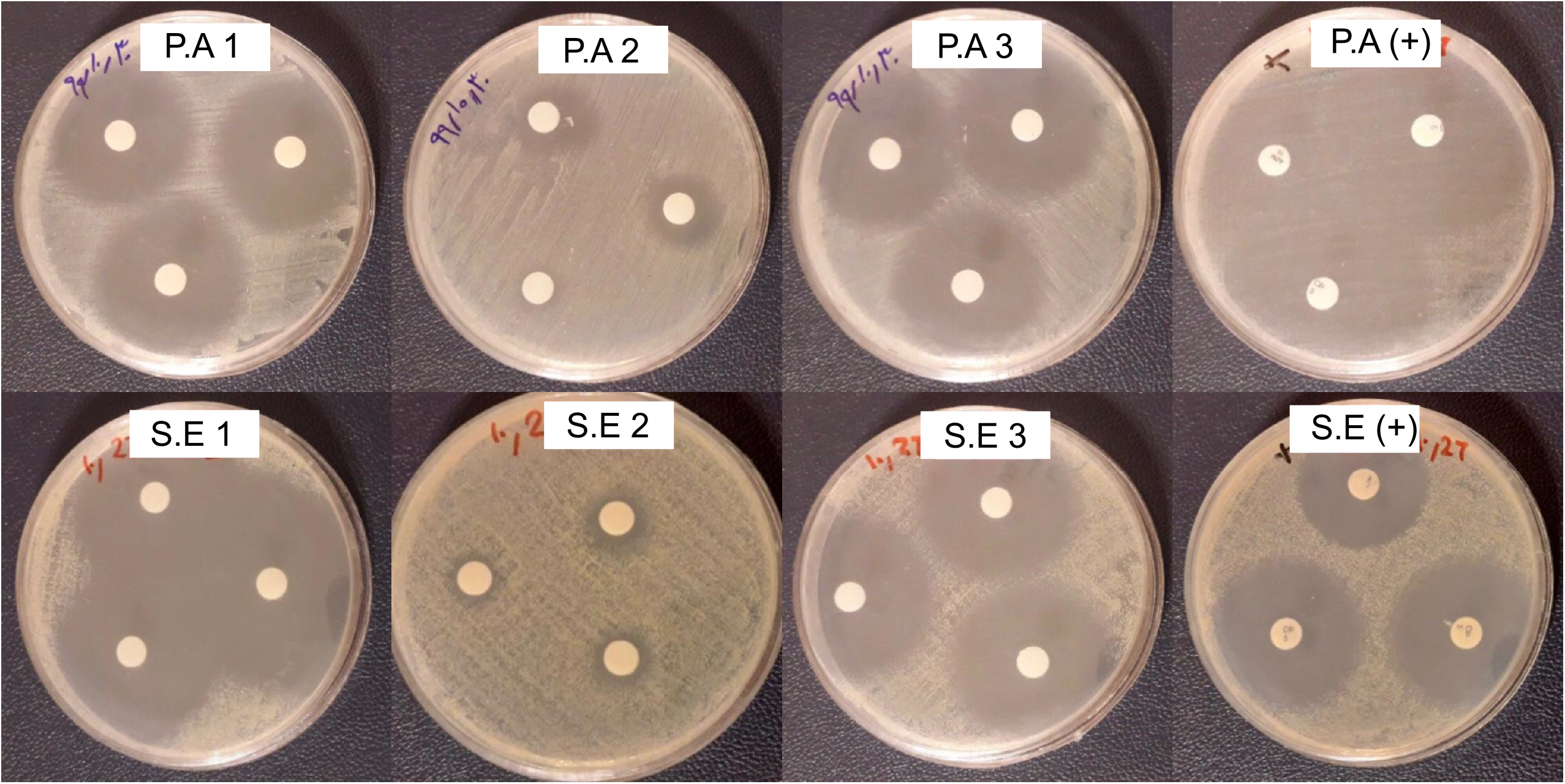
Growth-inhibiting against P.acne displayed with P.A 1, P.A 2, P.A 3, P.A (+) for optimal niosomal formulation, drug-free noisome, aqueous drug solution, and control sample, respectively. Furthermore, growth-inhibiting against S.epidermis displayed with S.E 1, S.E 2, S.E 3, S.E (+) for optimal niosomal formulation, drug-free noisome, aqueous drug solution, and control sample, respectively.

**Table 5.** The diameter of growth inhibition against P.acne and S.epidermis by applying N1 formulation, drug-free niosome, aqueous drug solution, and control (+) sample.

| Formulation | Inhibition zone (cm) |  |
| --- | --- | --- |
|  | P.acne | S.epidermis |
| N1 formulation | 2.76±0.05 | 3.66±0.05 |
| Drug-free niosome | 1.06±0.49 | 0.7667±0.12 |
| Aqueous drug solution | 2.63±0.05 | 3.233±0.05 |
| Control (+) | 3.66±0.05 | 3.46±0.05 |

### Conclusion

Four niosomal formulations with different percentages of their constituents (span 60 and cholesterol) containing doxycycline-hyclate were synthesized to treat acne. In this study, drug permeation through the rat skin occurred much faster than free drug when the niosomal formulation was applied, which indicated the dominance of the components of niosomal formulations over skin barriers and increased permeation of the drug through the skin. In evaluating in vitro drug deposition in rat skin, the optimal niosomal formulation with a 1: 1 molar ratio of span 60 / cholesterol showed drug deposition approximately three times higher than the control sample in the viable epidermis and dermis. The antibacterial activity of the optimal niosomal formulation, drug-free niosome, and free drug solution against the main acne-causing bacteria, namely Propionibacterium acnes and Staphylococcus epidermis, was evaluated by obtaining the diameter of the growth inhibition zone. The results of this evaluation showed the potential of doxycycline-containing niosomal carriers in antimicrobial activity. In addition, the cytotoxicity of the optimal niosomal formulation and free doxycycline on human dermal fibroblast cells was evaluated by the MTT method, which showed more excellent compatibility of the niosomal formulation for the cells. Overall, this study demonstrated the potential of an optimal niosomal formulation with a 1: 1 molar ratio of span 60 / cholesterol included doxycycline-hyclate, for acne treatment purposes.

## Acknowledgment

We thank Mr. Arya Aftab for his tireless help writing the article and presenting the language. We also thank Amirkabir University of Technology for providing the laboratory equipment and funding for this study.

## Author statement

The authors have no competing interests to declare.

